# Citizenship status and career self-efficacy: An intersectional study of biomedical trainees in the United States

**DOI:** 10.1101/2023.02.19.526477

**Authors:** Deepshikha Chatterjee, Ana T. Nogueira, Inge Wefes, Roger Chalkley, Susi Sturzenegger Varvayanis, Cynthia N. Fuhrmann, Janani Varadarajan, Gabrielle A. Jacob, Christiann H. Gaines, Nisan M. Hubbard, Sunita Chaudhary, Rebekah L. Layton

## Abstract

This study examines the intersectional role of citizenship and gender with career self-efficacy amongst 10,803 doctoral and postdoctoral trainees in US universities. These biomedical trainees completed surveys administered by 17 US institutions that participated in the National Institutes of Health Broadening Experiences in Scientific Training (NIH BEST) Programs. Findings indicate that career self-efficacy of non-citizen trainees is significantly lower than that of US citizen trainees. While lower career efficacy was observed in women compared with men, it was even lower for non-citizen female trainees. Results suggest that specific career interests may be related to career self-efficacy. Relative to US citizen trainees, both male and female non-citizen trainees showed higher interest in pursuing a career as an academic research investigator. In comparison with non-citizen female trainees and citizen trainees of all genders, non-citizen male trainees expressed the highest interest in research-intensive (and especially principal investigator) careers. The authors discuss potential causes for these results and offer recommendations for increasing trainee career self-efficacy which can be incorporated into graduate and postdoctoral training.

## Introduction

Biomedical programs in the United States (US) attract a significant number of international graduate and postdoctoral trainees. Since early 2010s, nearly 40% of PhD awardees in science and engineering were temporary visa holders (i.e., non-citizens; [1]). A significant percentage of these international trainees enter the biomedical workforce. Between 2014-2017, approximately 80% of visa holders intended to stay in the US, and about 45% of those with STEM degrees attained a job offer [2]. Similarly, the National Institutes for Health and Environmental Sciences reports that international trainees make up nearly half of their trainee population, of which about 45% stay in the US after completing their training (additional career outcomes data for specific institutions can be found through the Next Generation of Life Sciences Coalition transparent PhD career outcomes reporting initiative – e.g., UCSF). In fact, international trainees are crucial to the US COVID response [3] and to the US biomedical workforce broadly, with non-citizens and naturalized citizens making up 52% of the US biomedical workforce (as of 2014, compared with 21% in 1990;[4]).

Yet, non-citizen trainees in the US also face specific barriers [5] that are less understood by stakeholders in education and employment domains alike (e.g., academic mentors; funders; hiring, promotion, and tenure committees; and employers); furthermore, these systemic barriers are associated with deleterious career impacts [6–9]. In fact, the recent postdoctoral shortage, and parallel drops in international applications to biomedical doctoral programs, may be driven in part by international trainees’ reduced interest in seeking training in the US due to these barriers (7). In the current study, both doctoral and postdoctoral researchers were included as participants, and they are collectively referred to as “trainees.”

According to the social cognitive career theory (SCCT; [10,11]), one key variable that can mitigate the risk of negative career outcomes is career self-efficacy (CSE) [12], which is defined as the confidence with which people take charge of their career decisions. The SCCT describes how personal factors such as gender, race, socio-economic status, influence an individual’s career self-efficacy, which in turn influences career choices and outcomes. Self-efficacious people tend to have positive self-assessments about their capability to do well in a particular domain, such as in a chosen career. As such, they can readily adapt to various career demands and are also more proactive about managing their career choices [10,13, 14].

Biomedical non-citizen trainees tend to pursue research intensive or academic career paths [15]; based on the SCCT, it is possible that this pattern of career interests may be explained by their CSE which is examined in the current study.

In addition to assessing how trainees’ race and citizenship relate to CSE, the current study also examines how these factors relate to their pursuit of research-intensive PI-focused careers. Many trainees enter graduate and postdoctoral training with the intent to obtain research-focused faculty positions, although these interests wane over time (e.g., [16,17]). Multiple studies have sought to document and understand variations in these career preferences [16,18–24]). The few studies that include citizenship as a variable call for additional research (e.g., [16,25]). The impact of citizenship status remains understudied for biomedical trainees despite the fact that faculty career paths are often pursued by non-citizens, typically after postdoctoral training [26] (that may have occurred within or outside of the US). Importantly, the past decade has been marked by increasing awareness of valuing and preparing trainees for career paths across the scientific enterprise (beyond narrowly focusing on the faculty track) [27–33]. Nonetheless, the career interests of non-citizen trainees have received scant attention in the literature.

It is important to study the association between CSE and career interests in the context of citizenship because international trainees in the US face several systemic biases—both structural and interpersonal—that can negatively impact CSE. For example, citizenship status can constrain the types of job and learning opportunities in the US (detailed later). Furthermore, historically speaking, many immigrants to the US have often faced limited career opportunities even in their home countries. Non-citizen trainees’ CSE may be negatively impacted by factors, such as microaggressions, stereotyping, and biases in their daily work and career development [9].

According to SCCT, CSE relies on a constellation of factors, of which career socialization and the types of learning opportunities that are available to trainees are key influences [11,34,35]. In STEM fields, citizenship status and associated constraints seem to create systemic barriers that may negatively impact CSE; similarly, women and underrepresented racial/ethnic biomedical trainees face additional barriers than well-represented men [21]. The current study focuses on the intersection of gender and citizenship. However, race/ethnicity is not included because the categories that are commonly accepted in the US socio-political and historical context (e.g., Black, White, Hispanic) may not always be accepted by, or apply to, non-citizen trainees who come from different countries that might use different descriptors. In fact, for non-citizen trainees, national origin or cultural background could be far more salient components of identity than race and ethnicity [4,36]. Thus, to avoid forcibly lumping participants’ national and cultural heritage into US race-categories [37], the current study does not examine race/ethnicity.

Systemic barriers, such as US-centrism and patriarchal power structures, specifically restrictions on access to the US workforce due to complex visa regulations, significantly limit the opportunities of non-citizen trainees to develop evidence of their strengths and expertise tailored to their career interests (e.g., [15]. For example, Chatterjee [38] highlights how non-citizen trainees are often ineligible for prestigious federal fellowships and plum research projects (US citizenship or permanent residency is often a requirement). Some hiring committees penalize non-citizen candidates for the lack of US-only awards, grants, and other work experiences oblivious to the fact that these may have been inaccessible to non-citizen candidates.

In addition, visas can restrict where non-citizen trainees intern or work off-campus. The precarious position of visa holders was recently highlighted when, during the COVID-19 pandemic, the US government issued a policy that international students must leave the US if all their classes had been moved online [39]. This policy was later retracted for graduate students already in the US, but not for incoming students. Therefore, even before earning their doctoral degrees, structural limitations on training can restrict growth and developmental opportunities for non-citizen trainees. Similarly, postdoctoral scholars already training in the US were impacted by travel bans that made it difficult for them to exit/enter the US, and faced expensive and lengthy quarantines, new and renewal visa freezes, and subsequent processing delays due to backlogged cases [7]. US universities’ remarkable lack of investment in international students is evident, even though over the last two decades, depending on discipline, 32 to 43% of US Nobel Laureates [40] were foreign-born.

When trainees are not informed about such systemic barriers before they enter the US, and funders, employers, and others do not provide creative solutions to dismantle these barriers, then non-citizen trainees’ curriculum vitae can seem inferior relative to their citizen counterparts. Furthermore, the US limits the allocation of work visas in the private sector, thereby severely reducing the chance for equal employment opportunities for non-citizen trainees. Navigating attainment of a work visa can require high costs in both time and legal expenses—for trainee and/or employer—with high uncertainty of success; thus, some employers outright disqualify non-citizen job applicants. In addition to failing to provide structured information about these limitations, US institutions also fail to offer tailored training to mentors (who are predominantly US citizens) to offer customized advice to non-citizen trainees. As a result, non-citizen trainees are unable to make informed career decisions and are withheld from taking early steps to accrue the necessary experience and network that would help demonstrate their fit for different types of scientific roles. For example, Xu and colleagues [26] demonstrate that international postdoctoral fellows are more likely to stay in tenure-track academia or research-intensive roles, as opposed to US citizen counterparts—a finding mirrored in Mathur and colleagues’ [15] study for international graduate students whose top career outcomes included primary research type careers (in both academic and for-profit sectors).

The challenges and constraints faced by non-citizen trainees highlighted thus far are further compounded by individual identity characteristics, such as an individual’s gender, that can create additional barriers. Across cultures, women face additional systemic barriers of social, cultural, and familial expectations of marriage, child- and elder-care responsibility, and heavier expectations of contributing to household duties [41–44]. These burdens are further magnified in many international communities that may adhere to more traditional norms of what it means to be a woman, a mother, a wife, and a professional/academic, and how one must navigate these different roles at the same time [45,46]. The distinct experiences at the intersection of these identities (that is, being an international trainee and being female) should not be ignored. In ccordance with this view, intersectionality theory posits that it is important to consider how the different social identity groups to which people belong, taken together, impact key outcomes in many arenas [47]. Thus, the current study takes an intersectional perspective by looking at the joint effect of citizenship and gender on CSE and career choice.

The discussion so far highlights that CSE and career choice can both be contingent on factors such as gender and citizenship. Both these factors can constrain the career opportunities available to trainees. For example, women are typically undervalued in STEM fields by their mentors, peers, colleagues, and sometimes even their family members; such devaluation can challenge women trainees’ confidence and prevent them from choosing academic, research-intensive careers [48,49]. Even worse, these constraints may also influence their decisions to leave graduate and postdoctoral positions entirely [21]. Similarly, for non-citizen trainees, the burden of studying in a new culture, learning the norms and rules of a different educational setting than their home country, facing microaggressions, stereotypes and biases, concerns about language skills, accent and visa status-related barriers can all lower the trainees’ CSE and influence their daily work and career development [9,50–52] thereby constraining their career choices.

Taken together, and in line with the intersectionality approach, this study investigates the combined role of citizenship and gender on CSE. Belonging to two social identities that are traditionally seen as marginalized can have an enhanced detrimental impact on individuals than if they belonged to one marginalized social group only (known as double jeopardy). Because being a woman and being a non-citizen trainee are two social identities that are often viewed as marginalized, especially in the US STEM field context, it is likely that this double jeopardy is associated with lower CSE in female non-citizen trainees (compared to non-citizen males and compared to citizen males and women). This intersectional challenge is also likely to curtail career choices especially those that are already restricted for women non-citizen trainees.

## Research Aims

In sum, the current study uses data that had been collected by the NIH BEST survey of biomedical trainees with the aim to investigate two research questions: a) if citizenship (US citizen versus non-citizen) and gender (males versus females) impact CSE; and b) to what extent CSE is associated with specific career interests (e.g., pursuing careers as an academic or non-academic scientist. In doing so, the current study is responding to the recent calls by Ginther and colleagues [51] and Liu and colleagues [54] to assess the outcomes of those with intersectional identities in the biomedical field. The focus of this paper is to understand the differences in CSE and career choice for female non-citizen trainees, specifically considering the differential burden they may face compared to that of non-citizen male trainees, as well as in comparison to US citizen male and female trainees. Studies in other disciplines have just recently begun to explore CSE for doctoral trainees (e.g., [55]). The current study contributes to the biomedical workforce training development literature (e.g., [56]) with studies exploring factors that are associated with CSE of biomedical trainees (e.g., [25]), while adding an investigation of how citizenship status and gender together impact trainees’ CSE and career choice.

## Methods

### Participants

Participants were trainees (i.e., doctoral and postdoctoral) who were eligible to participate in NIH BEST programs, all of whom were invited to complete surveys (see [57]), from each of the 17 funded institutions nationally ([58] includes program highlights). Each institution either obtained approval of its own institutional review board or relied on the NIH-approved data-sharing agreement [57]. The current study is based upon common responses from standardized surveys across institutions. Respondents self-identified as male or female (choice did not indicate whether this referred to self-identified gender or biological sex; since the preferred term was self-selected hereafter the term gender is used, with the caveat that other gender options were unavailable for selection). Respondents also selected from the following citizenship categories: US citizens, temporary visa holders, and green card holders. For the analysis, participants with temporary visas or green card were classified as non-citizens, and those who reported being US or naturalized citizens were classified as US Citizens. This was done to reflect the systemic barriers that temporary visa holder non-citizens face. Furthermore, those with green cards—that is, expatriates and naturalized citizens—may still self-identify as international. Henceforth the terms “citizen” and “non-citizen” will be used (as opposed to international, while acknowledging that some of the barriers discussed may also be faced by internationally born US Citizens). A total of 10803 responses were collected. Some trainees did not disclose gender and citizenship data (n = 861). The present study includes 9942 participants, who identified collectively as citizens or naturalized citizens (n = 6077 citizens), and who identified collectively as temporary visa or green card holders (n = 3865 non-citizens).

**Table 1.**
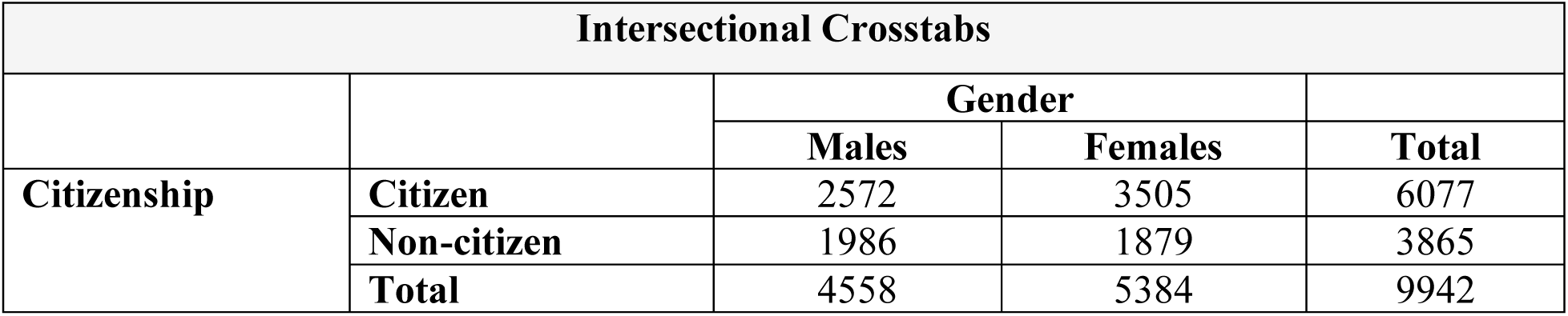
Number of respondents by citizenship and gender.

### Data Collection

The current dataset includes entrance surveys that were administered nationally during the first and second years of the NIH BEST cohorts (e.g., [57,59,60]). To maximize power and retain the largest sample of trainees, the current work is limited to entrance survey data only. To accommodate the variety of topics that the collective group of NIH BEST Consortium members requested to be addressed in the survey, study measures were developed by the NIH BEST contractor (Windrose) and harmonized by the consortium.

### Measures

*Career Self-Efficacy (CSE)* was assessed using a previously validated 5-item survey rated on a 5-point Likert Scale scale, from 1=not at all confident, 2=Minimally confident, 3=Moderately confident, 4=Highly confident, 5=Completely confident [59]. CSE was calculated as the mean of the following items for self-reported confidence to: “Assess your abilities to pursue your desired career path(s),” “Determine the steps to pursue your desired career path(s),” “Seek advice from professionals in your desired career path(s),” “Identify potential employers, firms, and institutions relevant to your desired career path(s),” and “Achieve your career goals” (Cronbach’s alpha = 0.86; see Supplemental Table 1 for complete analysis).

*Seniority* (years in the role as a doctoral or a postdoctoral trainee) was measured using data collected for the item Years-of-training).

*PI Career Interest* was assessed with the question, “to what extent are you currently considering [the career path of] Principal investigator in a research-intensive institution, using a 5-point scale, from 1=Not at all considering, 2=Slightly considering, 3=Moderately considering, 4=Strongly considering, 5=Will definitely pursue.” Similarly, the *Level of Consideration* was assessed for 20 common biomedical career pathway options (i.e., Principal Investigator (PI) in a research-intensive institution, Research in industry, Research staff in a research-intensive institution, Combined research and teaching careers, Teaching-intensive careers in academia, Science education for K-12, Science education for non-scientists, Clinical practice, Public health related, Scientific/medical testing, Science writing, Research administration, Science policy, Intellectual property, Business of science, Entrepreneurship, Sales and marketing of science-related products, Support of science-related products, Drug/device approval and production, Clinical research management. The variable *Sum of Career Paths Considered* for different career paths was assessed by combining the answers for each career path as “less interested” if equal or below 3 or “highly interested” if equal or above 4. The sum of interest was averaged and plotted on a 1 to 20 scale to evaluate how many career paths were being considered by participants.

The *Familiarity with 20 Career Paths* was measured on a 5-point Likert scale and as follows: 1=I am not familiar with any of these career paths, 2=I am familiar with a few of these career paths (between 1 and 6), 3=I am familiar with some of these career paths (between 7 and 12), 4=I am familiar with most of these career paths (between 13 and 19) and 5= I am familiar with all of these career paths.

*Career Training Attained* was assessed with the item, “I am **getting the training I need** for my desired career path(s),” hereafter *Career Training. Departmental Career Goal Support* was assessed with the item, “I am **encouraged** by my graduate program/department to pursue my career goals,” hereafter *Departmental Career Support*. Both used the 5-point Likert scale (0=Not applicable, 1=Strongly disagree, 2=Disagree, 3=Neutral, 4=Agree, and 5=Strongly agree).

*PI Encouragement* was assessed using a composite variable based on the mean score of two items, “I am encouraged by my PI/thesis advisor to **pursue career development activities** toward my career goals,” hereafter encouragement to pursue career development activities and “I am encouraged by my PI/thesis advisor to **pursue my career goals**,” hereafter encouragement to pursue career goals, using a 5-point Likert scale (0=I do not know, 1=Strongly disagree, 2=Disagree, 3=Neutral, 4=Agree, 5=Strongly agree).

### Analyses

Data analyses were completed using SPSS v27 (analysis of variance) and Graphpad Prism 9.4.1 (posthoc comparisons; figures), which were also used along with Illustrator 26.3.1 (figures) to produce data visualization. Only valid response options were included in the analysis. Posthoc Tukey corrections were used when relevant for multiple comparisons (Alpha = .05, two-tailed, between subjects t-tests).

First, one-way ANOVAS were used to test hypothesized effects of trainee status and seniority on mean CSE prior to establishing the final model. The remaining primary variables of interest were included in the final analysis of variance (citizenship status, gender, and PI career interest). Posthoc comparisons (independent samples t-tests) were conducted to identify intersectional interactions. Posthoc correlations were evaluated to identify relationships among key variables (Familiarity with 20 common biomedical career pathways; career training, departmental career support; PI encouragement) with career interest on CSE.

## Results

### Trainee Status and Seniority

Based on differences in career stage, it was hypothesized that trainee status (doctoral versus postdoctoral trainees) and seniority in each (years in trainee position) might be associated with career self-efficacy (CSE). However, neither of these showed significant association with CSE, and hence these variables were dropped from the final analysis (see Supplemental Table 2 for 4-Way factorial analysis of variance, including career status x gender x career interest x seniority).

Because no significant explanatory effects were identified, the simplified 3-Way ANOVA was retained for the final analysis.

### Intersectional Model: Factorial Analysis of Variance

A factorial Analysis of Variance (ANOVA) was conducted on the three variables of interest (career status x gender x career interest). The 3-way ANOVA was conducted to identify intersectional differences in CSE based on citizenship (citizen vs. non-citizen), gender (male vs. female), and career interest (PI vs. non-PI). All variables presented a significant main effect (see **Table 1**). P-values indicate the main effects for each variable of interest, \*\*\*\**p*<0.0001 and \**p*<0.05. **Figure 1** highlights the main effects of citizenship, gender, and career interest on career self-efficacy. As hypothesized, citizenship and gender showed a significant interaction (citizenship status x gender) on mean CSE. Contrasts were analyzed and Tukey-corrected for the variables citizenship x gender x career interest (**Supplemental Table 3**).

**Table 1.** 3-Way ANOVA: Citizenship, gender, and career interest.

| ANALYSIS OF VARIANCE |  |  |
| --- | --- | --- |
| Variable(s) | F-test | p-value |
| **Citizenship\*\*\*\*** | **50.918** | **< .0001** |
| **Gender\*\*\*\*** | **45.965** | **< .0001** |
| **PI Career\*\*\*\*** | **316.218** | **< .0001** |
| **Citizenship x Gender\*** | **6.264** | **.012** |
| Citizenship x PI Career | .242 | .623 |
| Gender x PI Career | .553 | .457 |
| Citizenship x Gender x PI Career | .216 | .642 |

**Figure 1a-c:**
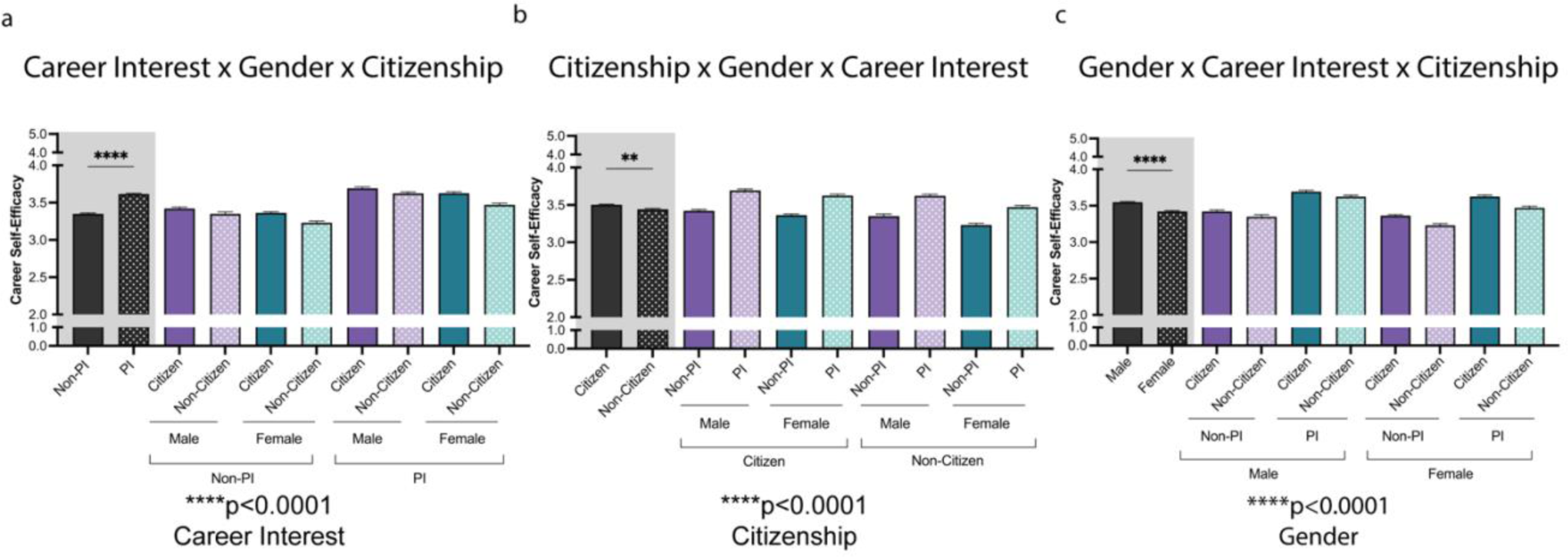
Visualization of Main Effects of Career Interest, Citizenship, & Gender on Trainee Career Self-Efficacy (CSE). A three-way ANOVA (citizenship, gender, PI career interest) showed a significant effect on CSE such that US citizens, males, and those with PI career interests were more efficacious. P-values at the bottom of each panel (a to c) indicate the main effects for each variable of interest, \*\*\*\**p*<0.0001. Post-hoc t-tests were conducted between the primary variable plotted on the left side of each panel (a-Non-PI vs. PI; b-Citizen vs. Non-Citizen; c-Male vs. Female); \*\*\*\**p*<0.0001and \*\**p*<0.01. Complete Tukey-corrected contrast analysis can be found in **Supplemental Table 3**. Black bars on the left side indicate the main variable. Purple and Green indicate the different variables: for 1a gender and citizenship, 1b gender and career interest, 1c career interest and citizenship.

Citizenship status impacted CSE such that non-citizen trainees (*M NC* = 3.44) were less self-efficacious than their citizen counterparts (*M C* = 3.50, *p*<0.0001) (see **Figure 2**). However, males were similarly self-efficacious regardless of citizenship status (*M C males*= 3.56; *M NC males*= 3.54). Females overall were less self-efficacious than males. Specifically, citizen females were more self-efficacious (*M C females*= 3.46) than non-citizen females (*M NC females* = 3.31).

**Figure 2:**
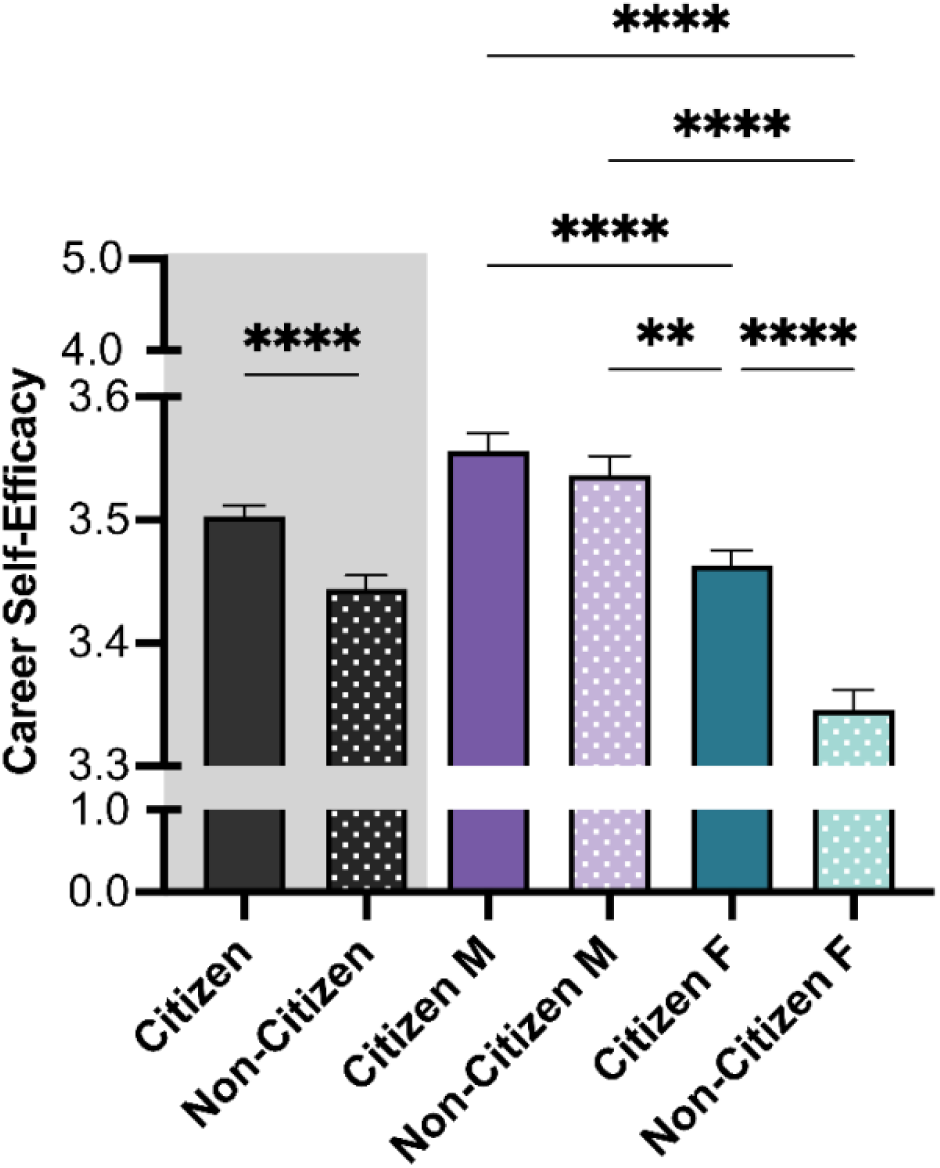
Main effects of Citizenship and Gender on trainee Career Self-Efficacy (CSE). Patterns for Citizenship and Gender showed a significant main effect, with higher CSE for those identified as citizen and male. Multiple comparison tests (Tukey-corrected posthoc t-tests) were corrected for with significant effects illustrated using brackets. P-values indicate significance of Tukey’s multiple comparison tests, \*\*\*\**p*<0.0001, and \*\**p*<0.01. *Note:* M=Male, F=Female. Color differences (purple and green) indicate the main variable gender, where purple = male, green = female; pattern indicates citizenship (no pattern = citizen; pattern = non-citizen).

### Differences Between International and Domestic Trainees in Pursuing Potential Career Paths

We next analyzed the interest in pursuing a PI career at a research-intensive institution. The strongest effect was detected between non-citizens versus citizen trainees, as displayed in **Figure 3**. Non-citizen trainees were most interested in pursuing a PI career (*M NC* = 3.42, *M C* = 2.97). Relative to their citizen counterparts, both non-citizen men (*M NC* males = 3.72 vs *M C Males* = 3.21) and women (*M NC* females= 3.10 vs. *M C* females= 2.79) showed higher interest in pursuing a PI career. Of note, citizen women presented the lowest interest.

**Figure 3:**
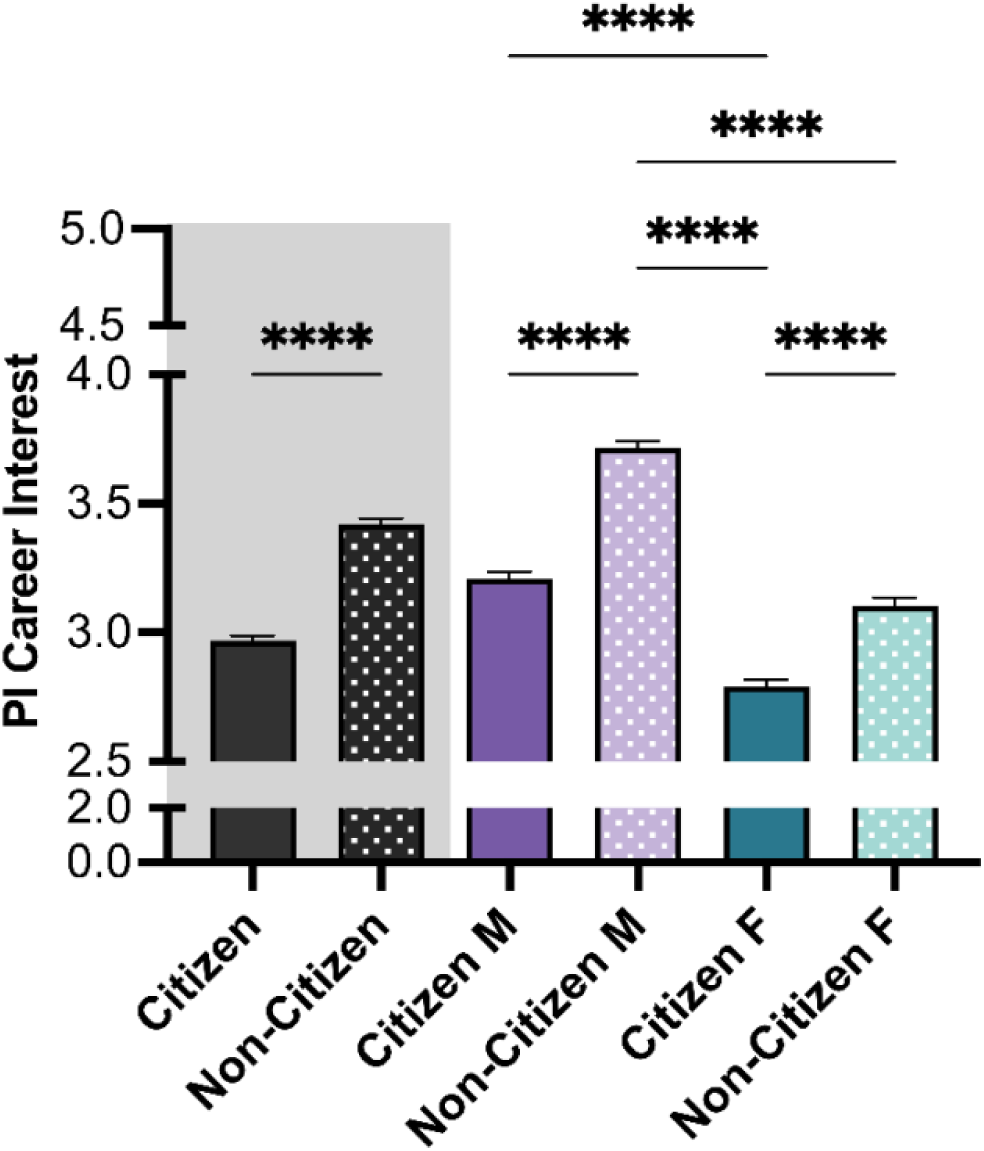
Confidence in pursuing a PI Career Path. Citizenship and Gender showed a significant main effect in the interest to pursue a PI career path, with highest confidence for those identified as non-citizen and male. Post-hoc t-tests were conducted between all possible pairings, as illustrated by each end of the respective bracket. P-values indicate significance of Tukey’s multiple comparison tests, \*\*\*\**p*<0.0001. *Note:* M=Male, F=Female. Black bars indicate the main variable citizenship. Color differences (purple and green) indicate the main variable gender, where purple = male, green = female; pattern indicates citizenship (no pattern = citizen; pattern = non-citizen).

Next, career paths considered by both citizen and non-citizen trainees were evaluated (**Figure 4**). Both citizen and non-citizen trainees showed more interest (M > 2.7) in non-leadership research-intensive positions, including research in industry. Accordingly, non-citizens and citizens showed less overall interest in other career paths (M < 2.3), including teaching-intensive and research-adjacent career paths (Supplemental Figures 1 & 2, respectively). The low interest in all career paths except research was striking (Supplemental Figures 1 to 4).

**Figure 4:**
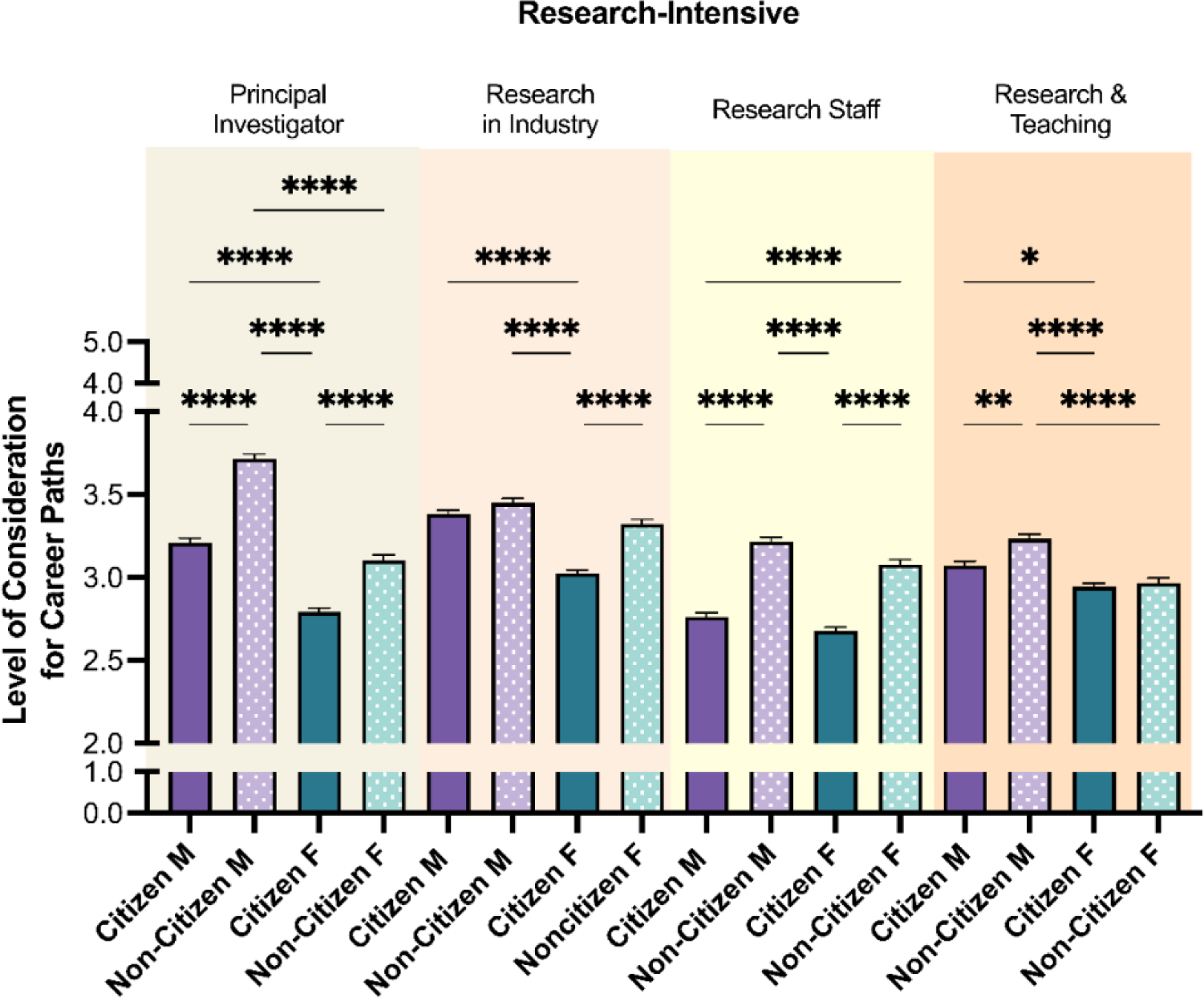
Level of consideration for research-intensive career paths. Post-hoc t-tests were conducted between all possible pairings within each career path, as illustrated by each end of the respective brackets. P-values indicate significance of Tukey’s multiple comparison tests, \*\*\*\**p*<0.0001, \*\**p*<0.01, and \**p*<0.05. *Note:* M=Male, F=Female. Color differences (purple and green) indicate the main variable gender, where purple = male, green = female; pattern indicates citizenship (no pattern = citizen; pattern = non-citizen).

Although both citizen and non-citizen trainees scored higher interest in research-intensive careers (**Figure 4**), the average number of careers considered by trainee groups differed (**Figure 5**). Non-citizen trainees contemplated more career paths than citizen trainees (*M NC* = 3.86, *M C* = 3.17). Non-citizen men showed the highest number of various career interests (*M* NC *males*= 4.11,) followed by non-citizen women (*M NC females*= 3.60). The lowest number of various career interests were selected by citizen females (*M C females* = 3.03). Of note, amongst both citizen and non-citizen trainees, there was a high interest in research in industry with non-citizen men and women having the highest interest compared to their citizen counterparts.

**Figure 5:**
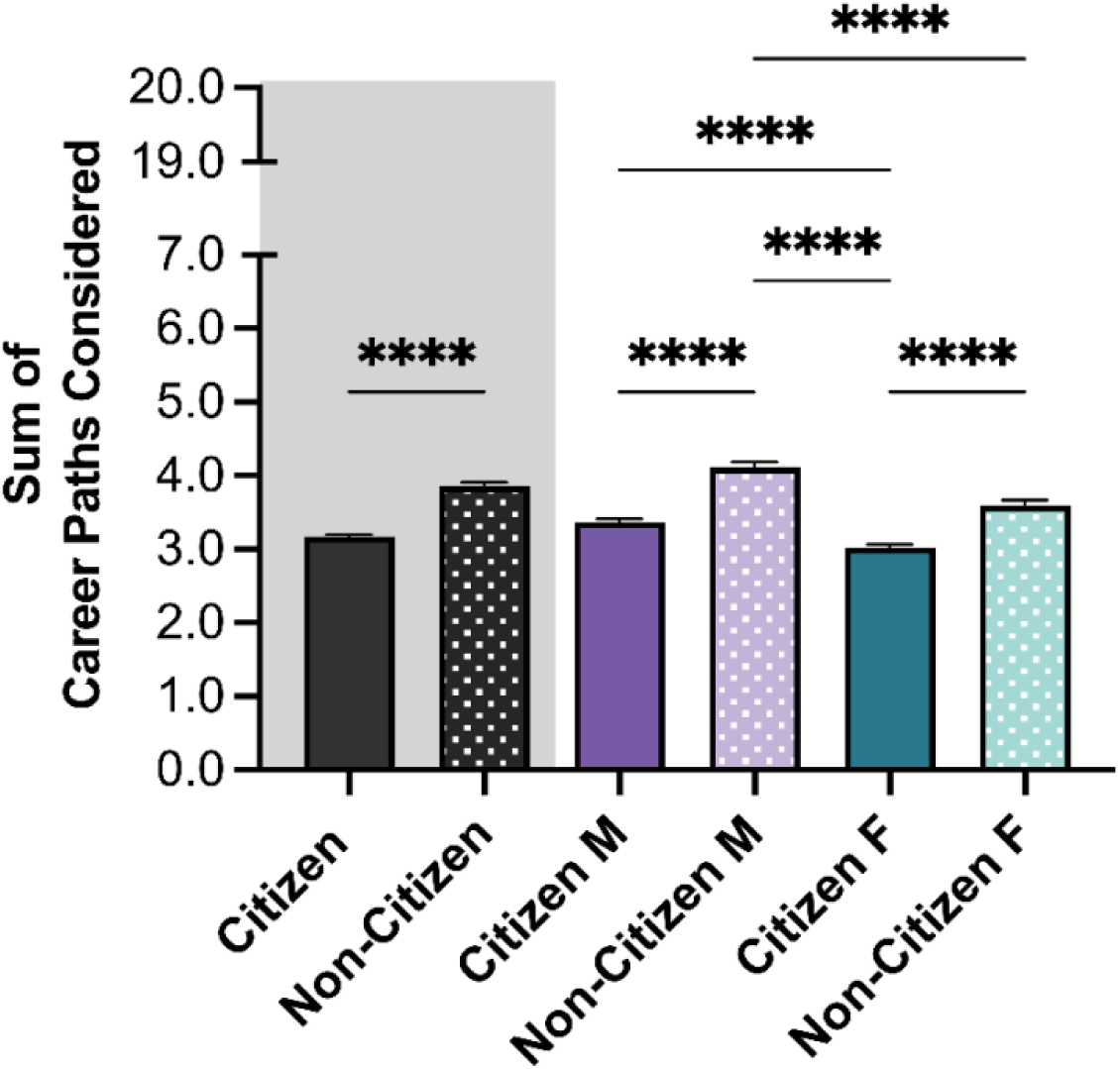
Career Path Interests. Career Path Interests was calculated as the sum of career paths considered (up to 20 items, could be selected). Citizenship and gender showed a significant main effect on the number of career paths considered by trainees. See Methods for details about the creation of Sum of Career Paths Considered. Post-hoc t-tests were conducted between all possible pairings, as illustrated by each end of the respective bracket. P-values indicate significance of Tukey’s multiple comparison tests, \*\*\*\**p*<0.0001. *Note:* M=Male, F=Female. Color differences (purple and green) indicate the main variable gender, where purple = male, green = female; pattern indicates citizenship (no pattern = citizen; pattern = non-citizen).

Further investigation of the trainees’ knowledge of 20 common biomedical career pathways revealed that non-citizen trainees reported less familiarity with the different career options (*M NC* = 3.34, *M C* = 3.88) (**Figure 6**). Interestingly, non-citizen female trainees had more familiarity than non-citizen male trainees (*M NC female*= 3.40, *M NC male*= 3.29). No difference was detected between male and female trainees among citizens (*M C female*= 3.86, *M C male*= 3.89). Pearson correlation coefficient was used as a post-hoc analysis to investigate the relationship between familiarity with the different career paths and level of CSE. There was moderate-positive, but significant relationship between these two variables, such that the more informed trainees were about the different career paths, the more career efficaciousness they reported (*r* = 0.231, *p* < 0.001).

**Figure 6:**
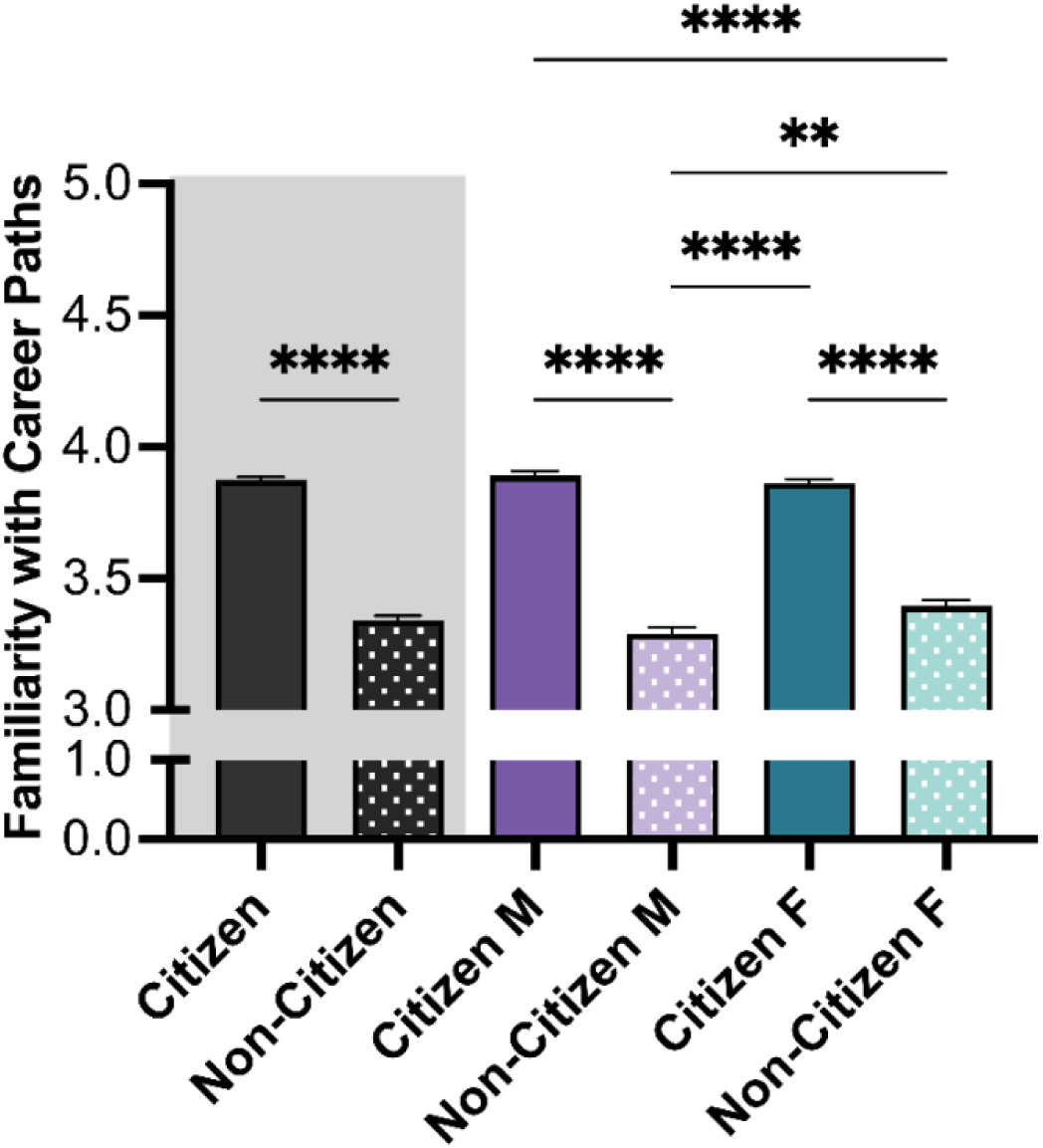
Familiarity with 20 common biomedical career pathways. Citizenship and Gender showed a significant main effect on the familiarity with different career paths, with higher confidence for those identified as citizens. Post-hoc t-tests were conducted between all possible pairings, as illustrated by each end of the respective bracket. P-values indicate significance of Tukey’s multiple comparison tests, \*\*\*\**p*<0.0001 and \*\**p*<0.01. *Note:* M=Male, F=Female. Black bars indicate the main variable citizenship. Color differences (purple and green) indicate the main variable gender, such that purple = male, green = female; pattern indicates citizenship (no pattern = citizen; pattern = non-citizen).

### Training and Encouragement to Pursue Desired Career

Using the data collected, the training needed to pursue desired career path (Figure 7, Career training attained) and the level of encouragement reported from the graduate program (for doctoral trainees) or from the department (for postdoctoral trainees) (**Figure 7**, Departmental Career Goal Support) were investigated. There was no significant difference between citizen or non-citizen trainees as both reported feeling neutral about needing more career-related education to pursue their career goals (*M C* = 3.63, *M NC* = 3.67) (**Figure 7**, Career Training Attained). This trend was similar when the intersectionality of citizenship and gender was analyzed, with the exception of non-citizen male trainees, who reported feeling more strongly about having obtained the requisite training in comparison to other groups. Similarly, trainees reported feeling neutral about levels of support from their graduate program/department in helping them to pursue their career goals. However, non-citizen trainees reported feeling less support from the graduate program/department than citizen trainees (*M NC* = 3.47, *M C* = 3.72) (**Figure 7**, Departmental Career Goal Support). There was no difference when assessing the intersectionality of citizenship and gender and the level of support reported by trainees.

**Figure 7:**
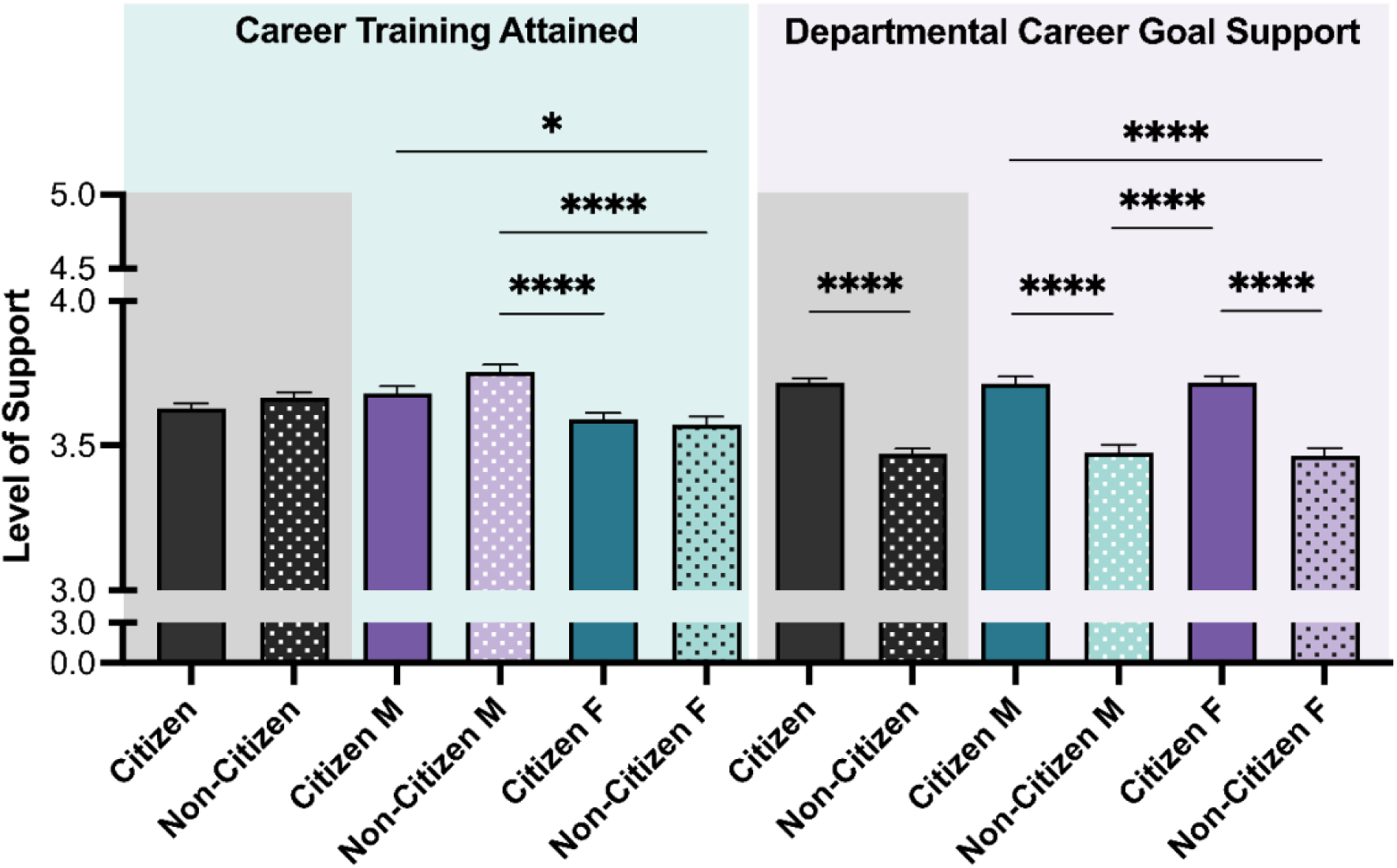
Career Training Attained and Departmental Career Goals Support ratings. The level of support reported by trainees in pursuing their desired career path was evaluated with two items: Career training attained and departmental career goal support. Post-hoc t-tests were conducted between all possible pairings, as illustrated by each end of the respective bracket. P-values indicate significance of Tukey’s multiple comparison tests, \*\*\*\**p*<0.0001 and \*\**p*<0.01. *Note:* M=Male, F=Female. Black bars indicate the main variable citizenship. Color differences (purple and green) indicate the variable gender, where purple = male, green = female; pattern indicates citizenship (no pattern = citizen; pattern = non-citizen).

To assess the association between obtaining sufficient training and level of CSE a Pearson correlation coefficient was computed. Not surprisingly, there was a significant positive relationship between these two variables, as the more training that was acquired, the more career self-efficaciousness trainees reported (*r* = 0.323, *p* < 0.001). Similarly, when trainees reported feeling more supported by the graduate program/department, they also reported having higher career self-efficacy (*r* = 0.286, *p* < 0.001).

The level of encouragement by their PIs to pursue career development activities and career goals reported by trainees was evaluated (**Figure 8**). Since the question and trend were similar, these two questions were combined into a single variable, which was labeled PI encouragement. The encouragement reported by non-citizen trainees was higher in comparison to citizen trainees (*M* N *C* = 3.33, *M C* = 3.19). This trend was similar between non-citizen and citizen male trainees (*M NC males* = 3.41, *M C males* = 3.19). There was no significant difference in the level of encouragement reported between non-citizen and citizen female trainees (*M NC females* = 3.19, *M C females* = 3.26) (**Figure 8**). Post-hoc analysis revealed that the level of encouragement trainees experienced from their PI was moderately related to CSE. A significant positive relationship was found between these two variables, such that higher PI support was associated with higher CSE (*r* = 0.182, *p* < 0.001).

**Figure 8:**
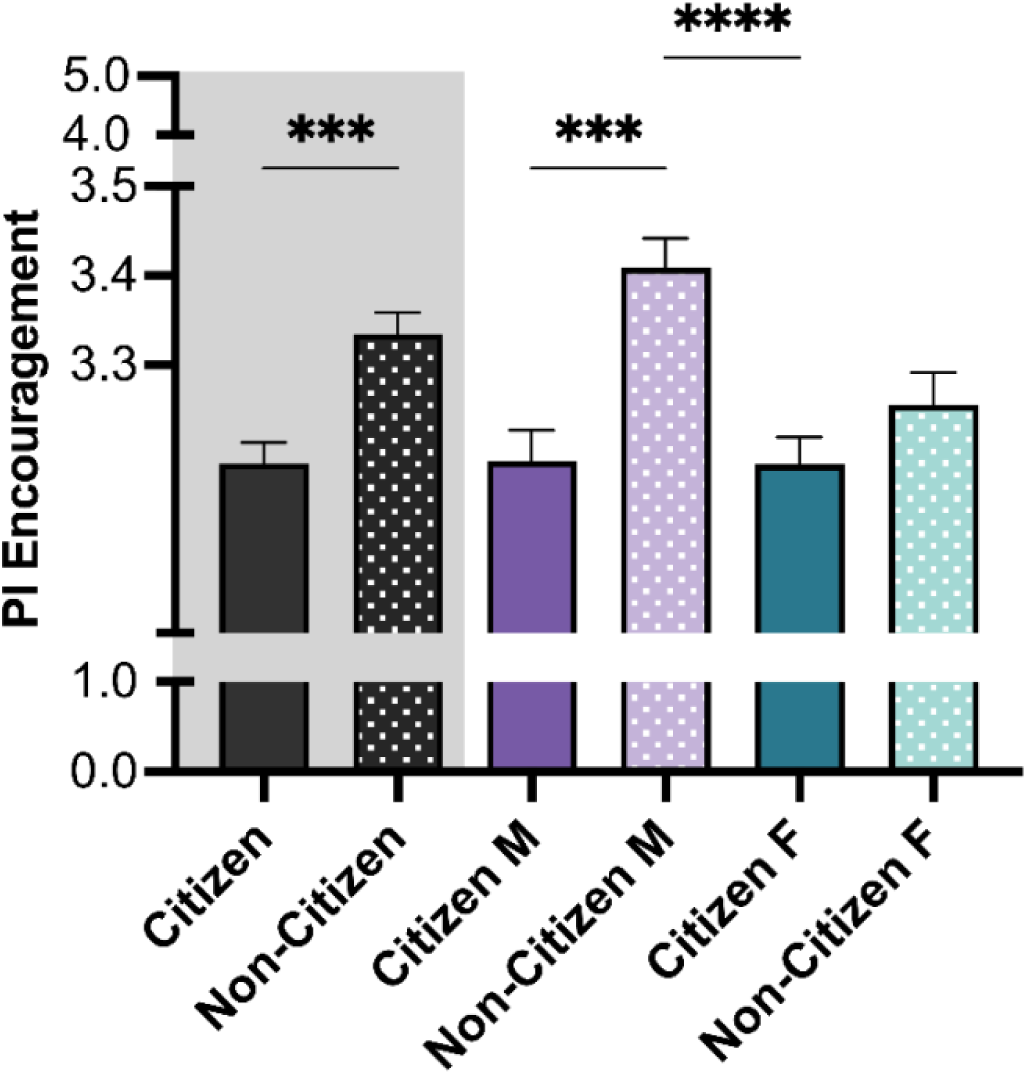
PI Encouragement reported by trainees. PI encouragement was measured combining two items: Encouragement to pursue career development activities, and encouragement to pursue career goals. Post-hoc t-tests were conducted between all possible pairings, as illustrated by each end of the respective bracket. P-values indicate the significance of Tukey’s multiple comparison tests, ****p<0.0001 and ***p<0.001. *Note:* M=Male, F=Female. Black bars indicate the main variable citizenship. Color differences (purple and green) indicate the variable gender, where purple = male and green = female; pattern indicates citizenship (no pattern = citizen; pattern = non-citizen).

## Discussion

The present study was undertaken in response to calls for studying intersectional effects on career outcomes [53,54]. It included citizenship as a key variable because research on career outcomes for non-citizens in the US remained sparse despite STEM fields attracting trainees from all over the world for doctoral and postdoctoral work. To address this gap, the current study had two key goals: (a) to assess how citizenship and gender relate to career self-efficacy (CSE) of doctoral and postdoctoral trainees, and (b) to assess if interest in PI/research intensive academic careers is related to CSE. CSE was defined as the confidence with which people take charge of their career decisions and was measured using a validated CSE scale [59]. To a large extent, studies on the diversity of trainees in STEM fields have been conducted with undergraduates whereas research at the doctoral and postdoctoral level is sorely needed given the unique stressors, challenges, and barriers that these environments present for trainees.

For the first research aim of assessing intersectional effects of gender and citizenship, the current study found that overall females reported lower CSE than males; this is in line with previous literature that has examined these effects in the biomedical context [21,61].

Furthermore, non-citizen trainees had lower CSE than citizen trainees. When the intersectionality of citizenship and gender on CSE was assessed, it was found that there was a double disadvantage to non-citizen women as their CSE was lowest (compared to that of non-citizen men, citizen men, and citizen women). The disparity in self-efficacy for non-citizens (in particular non-citizen females), is concerning. This outcome supports the double jeopardy hypothesis, such that for people who hold two socially marginalized identities, the negative effects are exacerbated; the seniority of the trainees did not affect this outcome. Future research should focus on intersectionality to better understand how different configurations of identities could be marginalized in STEM fields (e.g., studying varying combinations with race/ethnicity, citizenship status, neurodiversity attributes, gender, and other identities in an intersectional manner). Studying combinations of different social identities holistically to understand the heterogeneity inherent within different groups is also vital.

For the second research aim, the current study investigated how one’s interest in pursuing an academic PI/research intensive career related to CSE. Remarkably, while overall, non-citizen trainees had lower CSE than citizen trainees, they nonetheless showed higher interest in PI careers than their citizen counterparts. Similarly, previous work, though not accounting for differences by citizenship status, found that fewer female postdoctoral trainees showed interest in pursuing PI career (e.g., [62]). Martinez and colleagues (2007) offered family considerations/commitments (especially childcare), and lower confidence in succeeding in a PI career path as primary reasons for this difference. Additionally, a majority of female biomedical trainees’ intent on pursuing PI careers expressed facing gender-conscious experiences and strategized managing their identities [63].

It may be possible that non-citizen trainees originally pursued training in the US with the belief that there were more opportunities to obtain a PI position relative to their country of origin. However, the wide variety of visa restrictions during training and post-graduation may have limited their career choices and interests. Non-citizen trainees may present lower CSE as they are ineligible to participate in prestigious research programs (e.g., federal funding such as NIH K99-R00, F-31, F-32 and NSF fellowships are available only for citizens), and have reduced ability to participate in external industry-focused roles due to visa restrictions, each of which could be viewed as a “golden ticket” for some to a prosperous career [38, 64]. These restrictions can also withhold them from working in start-ups [65]. Such policies and restrictions can influence trainees to opt for academic roles in order to maintain or retain their residency status in the US.

Finally, the current study highlights how CSE at the doctoral/postdoctoral levels remains low overall. This is a concerning finding and merits consideration in future policies, not the least because it could contribute to the mental health crisis in this population. This could be due to the scientific training process that can feel scathing as trainees are exposed to threatening negative feedback from their PIs, reviewers, and funders more often than positive mentoring messages.

Many doctoral and postdoctoral training standards are still geared toward preparing trainees for such roles. In line with the SCCT, trainees’ own normative expectations of what the graduate education enterprise entails can be strengthened when they receive congruent messages from their surroundings (i.e., their departments, faculty mentors, peers). In other words, the alignment (or misalignment) between one’s self-beliefs and the values that are communicated by the environment can together create powerful mechanisms that enhance (or reduce) CSE, especially for those interested in academic, faculty, and PI-careers.

Future research may be needed to explore insights highlighted in the posthoc analysis. Access to training opportunities, level of support reported at the departmental level, and PI encouragement potentially affect trainees’ self-efficacy. Both non-citizens and citizens reported that career training met their needs (career training attainment) (Figure 7). Interestingly, however, the encouragement reported by non-citizens at the department level was significantly lower than the encouragement reported by citizens (Figure 7). In contrast, citizens reported less encouragement by their PI than non-citizens (particularly non-citizen males who reported the most encouragement). This may be due to the higher interest of non-citizens (particularly non-citizen males) in PI careers (Figure 4) who may feel more encouraged within the academic career setting. Trainee career advancement is intrinsically connected to their relationship with their PIs [66–70]. Although it obvious that is crucial for those pursuing a PI or faculty career track, PI support is important for all trainees. One take-away may be that PIs could be more cognizant about providing encouragement for students interested in other career paths. Together these findings suggest citizenship status may pose potential barriers to integrating with colleagues in the lab and at the adjacent spaces. Lack of familiarity with different career options along with other barriers, such as visa restrictions, and the scarcity of academic positions, could impair the preparedness of non-citizens for long-term career outcomes.

Concurring with previous research showing that international trainees prefer PI careers [16], it is important to note that in the current study, although PI and research-intensive career interests were most prevalent for non-citizens, some non-citizen trainees are interested in other career paths than PI/academic research-intensive careers. In fact, they show interest in a wider variety of career paths than citizen trainees. Non-citizen males were interested in a higher number of careers relative to non-citizen females. It is difficult to assess why international trainees lack familiarity with the different career paths. Perhaps it is the case that non-citizen trainees, compared to citizens, over time learn about the visa restrictions and these constraints make PI career their most viable choice; alternatively, it could also be the case that they consider these positions to be more prestigious opportunities. However, the possibility of the latter alternative explanation is lower since academic jobs have lost their luster in most disciplines including biomedical science (e.g., national coverage on the state of tenure-track jobs in biomed; the Great Resignation and #leavingacademia trend; reports on job losses and salary cuts) [71–73]. Even when there are opportunities to engage in more career exploration, it is likely that non-citizen trainees simply devote their time to other research-relevant activities. In fact, post-hoc findings seem to support this idea because even when non-citizen trainees consider different career paths, they still prefer careers that are research-intensive non-academic paths.

Taken together, the post-hoc findings described above suggest that non-citizens may feel pressured into PI roles due to their precarity in the US as non-citizens. Future research should assess if non-citizen trainees’ lack of familiarity might be passive (i.e., something that happens due to lack of opportunities to gain exposure to different careers) or if they are making an active choice (i.e., prefer focusing on roles similar to their current positions and training background). Furthermore, investigating whether the escalation of commitment to becoming a PI and/or being on a research-intensive career track may be influenced by perceptions of precarity due to visa status, sunk costs (e.g.,[74]), reactions to cognitive dissonance [75], familial pressure for the prestige of research or academia in their home countries (e.g., losing face in collectivist cultures; [76]), or other reasons.

Another explanation for these findings may be that non-citizen trainees may not know that it is acceptable to ask about career training opportunities/resources that might help with their career interests and goals. This discomfort could stem from a myriad of reasons, including uncomfortable interpersonal interactions (e.g., experience of microaggressions). Indeed, due to cultural norms toward high power distance some trainees may feel it is improper to ask for help unless offered (a feeling of being a guest and not wanting to overstep their welcome) or may just be concerned about coming across as dependent on their advisors. If they also happen to be first-generation students, then hidden curriculum (e.g., the unspoken rules and cultural expectations of educational institutions; [23,77]) could make trainees’ hesitant to seek career training support. Potential fear of a negative impact on how they are viewed, fear of negative repercussions of asking, fear of being taken off choice projects if they reveal alternative career interests or need for the PI’s sponsorship of their visa or immigration applications may be additional concerns. These concerns can be further exacerbated by the power differential that arises from their reliance on funding of their work by their PI versus feelings of entitlement when they are “free to their lab/PI”, (i.e., on extramural funding such as supplements or fellowships).

### Implications and recommendations

First, the issue of low CSE at the predoctoral and postdoctoral levels needs to be addressed by creating systemic solutions. The analyses presented also show that all biomedical trainees— regardless of citizenship and gender—need to be better supported to strengthen their CSE. Furthermore, previous literature shows there is a need for development of systemic solutions to better support biomedical faculty so they can effectively advocate for biomedical trainees [31]. Understanding the reasons why CSE remains low is a vital next step. Once these reasons are elucidated, figuring out the mechanisms that could mitigate low confidence would need to be explored in greater detail, especially accounting for the intersectionality of trainees’ identities.

Second, understanding that not all non-citizens are alike will be important. Faculty mentors, graduate program directors and administrators, industry stakeholders all need to develop cross-cultural competence and communication skills that at the very least attempt to be respectful about, if not tailored to, non-citizens’ preferences to promote CSE. This attention is especially relevant to those who serve on admissions and hiring committees. Being knowledgeable and mindful about the type of implicit and silent constraints and systemic barriers that non-citizens encounter would be a great first start. It is recommended that funding agencies, universities and graduate programs all create focused programs of offerings that shed light on these issues. Especially in the United States, where the immigration pathway is broken for many (e.g., Indian and Chinese nationals for whom the path to green cards can take decades), and hardly ideal for other nationalities, it is surprising that these barriers remain virtually unknown, unconsidered, and ignored despite the fact that in STEM fields there is no paucity of international talent. It is time to ask why this is the case, and how this can be remedied.

Third, encouraging help-seeking behavior and creating inclusive professional development programs that specifically support non-citizens and women trainees are needed. For instance, Optical Practical Training (OPT) for international graduate students has been one essential resource to gain professional development and critical experience prior to beginning their graduate or post-doctoral work [78]. However, OPT is expensive, limited to very a specific type of visa (F-1 Student Visas), and applying and receiving approval for this extension is not a guarantee. Furthermore, this option is typically not available to postdoctoral trainees and is only temporary in nature. This leaves many potential trainees without the experience that their US counterparts have access to. Therefore, there is a need for departments/institutions to provide customized programming, support, and encouragement for trainees to explore and make informed choices about their career (i.e., addressing grant, program and visa limitations. In addition, establishing a mentoring program or mentoring committee that focuses on career advancement (whether in PI or non-PI roles), especially targeted towards non-citizens and women trainees could help support those with high interest but CSE to succeed in their desired career roles.

### Limitations & Future Directions

Although the primary analysis of this paper was to address issues impacting international trainees, it is important to note, the NIH BEST survey was not tailored to assess the experiences and/or concerns of non-citizen trainees. Survey questions relevant to understanding their current challenges and progression in doctoral and postdoctoral work were lacking. For example, the survey did not ask about country of origin, and instead only asked about current (US) citizenship status. Similarly, questions on how long trainees had been in the US, and for postdocs if they obtained their doctoral training in the US, would have been helpful to better understand adaptation and acculturation. While time in graduate school or postdoctoral training could be used as a proxy for at least a minimum time spent in the US, the time factor could not be assessed directly based on the current data set. Similarly, since self-concepts evolve over time, it is hard to say at what point a foreign national who emigrates to the US and becomes a US citizen self-identifies as such. However, the authors chose to focus on examining the impact of barriers faced disproportionately by those without US citizenship. Future work should study the length of stay in addition to other factors that impact trainees’ adaptation, acculturation, and/or assimilation to the host culture, which could in turn enhance career self-efficacy of biomedical graduate and postdoctoral trainees. It was assumed that the career goals for citizens and non-citizens would be similar when, in fact, the career concerns of non-citizens could be qualitatively different, especially as they relate to pursuing training for long periods in the US while away from family and home country. Questions about their understanding of the work visas/green cards after education were not asked, so it is unclear if non-citizens viewed promising career pathways as those that would lend themselves to permanent residency and/or future citizenship applications; if they would like to work in their home countries; or explore other global opportunities. Future research may include the impact of geographical contextual variables, such as country of origin, desired work location, and desired citizenship status on CSE and career outcomes.

Another limitation was that non-citizen trainee populations are quite heterogeneous. They come from a diverse set of cultures, espouse different values, and their educational journeys in higher education in the US may therefore take different forms impacting career path familiarity, how they envision their careers, their career goals and how they pursue them. Furthermore, the current study cannot account for any culture and/or national origin related differences in perceived or actual barriers that would impact CSE or career interest. For example, Indian and Chinese PhD holders, compared to trainees from other countries, face a particularly long and uncertain path to working in industry and to becoming a citizen in the US. Even after being eligible for applying for green cards, their green card application process can take decades whereas for trainees from other parts of the world it takes anywhere between 2-3 years. Similarly, trainees who want to go back to their host countries may face these and other barriers impacting their career outcomes [26] (Xu et al., 2018). Even with these limitations, the current study is important because it is one of the first studies to use an intersectional lens to study the impact of citizenship and gender, and the findings show that citizenship is a key variable for self-efficacy that must be examined in future work, ideally comparing international student and postdocs from different cultures.

Just as non-citizen trainees were not explicitly considered in the design of the BEST Survey, neither were female (or gender-non-binary) trainees. Even though there is research that shows that women must often navigate work-family conflict, and encounter spillover effects of family-to-work and work-to-family [79], there were no questions about these aspects in the current survey. These burdens would also apply to women who are US citizens but are likely even more pernicious to non-citizen women as they must contend with greater gender-identified cultural demands and roles (e.g., homemaker, or high expectations that a good career is one that does not take much time away from family). Next steps for future research may include a more formal evaluation of issues such as care-giving burden, work-life conflict, and other issues disproportionately faced by non-citizen female trainees. Furthermore, future research should include multiple gender identities to better understand unique challenges faced by trainees identifying as such.

The current study is also a call to both academia and industry to actively invest relevant resources to better understand and address the barriers faced by the international trainee workforce, including consideration of funding mechanisms for support [80]. It is vital that programs focused on diversity, equity, and inclusion, as well as assessments from graduate schools, postdoctoral offices, policymakers, funding agencies, and others, include non-citizen trainees and women as key stakeholder groups.

## Conclusions

Career self-efficacy was lower for non-citizen trainees compared to their citizen counterparts, and the lowest for non-citizen female trainees. Career interest differences between international and citizen trainees may be due to self-selection toward traveling abroad and research-intensive careers (due to visa requirements, high activation energy to be competitive for visa, and high cost of visa applications). Although non-citizen trainees expressed more interest in different career opportunities, ultimately, they showed lower familiarity with career options. It is unclear if non-citizen trainees have fewer options for career development training while facing a new training environment, or if they are not interested in such information and choose not to attend if they do not feel it is relevant to their goals. Non-citizen trainees demonstrate higher interest in different career opportunities yet remain loyal to PI careers; this may point to visa-related precarity as a driver of constrained career choices. Non-citizen trainees seem to have a stronger interest in academic PI-careers but it is unclear if this is due to personal career interests or to protect themselves from the systemic visa barriers. Future research should include a more in-depth analysis of the needs of non-citizen and female trainees’ pursuit of career development opportunities and their unique challenges.

