## Supplemental Files for "Citizenship status and career self-efficacy: An intersectional study of biomedical trainees in the United States"

**S1 Table. Summary of study variables, key constructs, common abbreviations, & survey questions**

| Variable name | Transformed | Survey Question(s) Used | Scale | Notes & Abbreviations |
|---|---|---|---|---|
| Career Self-Efficacy | Bivariate | Assess your abilities to pursue your desired career path(s)<br>Determine the steps to pursue your desired career path(s)<br>Seek advice from professionals in your desired career path(s)<br>Identify potential employers, firms, and institutions relevant to your desired career path(s)<br>Achieve your career goals | 1=Not at all confident<br>2=Minimally confident<br>3=Moderately confident<br>4=Highly confident<br>5=Completely confident | SCCT=Social Cognitive Career Theory<br><br>CSE= Career Self-Efficacy |
| Seniority | Bivariate | Years in current position | Junior vs. Senior | Graduate Student: Junior = 3rd year and below; Senior = 4th Year and up<br><br>Postdoc: Junior = 1st year and below; Senior = 2nd year and up |
| Gender | N/A | What is your gender? (optional) | 1=Male<br>2=Female | |
| PI Career Interest | Bivariate | To what extent are you currently considering [the career path of] Principal investigator in a research-intensive institution | 1=Not at all considering<br>2=Slightly considering<br>3=Moderately considering<br>4=Strongly considering<br>5=Will definitely pursue | PI=Principal Investigator (research-intensive)<br>a) If equal or below 3 = Non-PI;<br>if equal or above 4 = PI<br>b) The biomedical career pathways listed were coded as described for PI Career Interest |
| Sum of Career Path Considered | Numerical | To what extent are you currently considering [the career path of] [INSERT CAREER PATHWAY TITLE] | 1=Not at all considering<br>2=Slightly considering<br>3=Moderately considering<br>4=Strongly considering<br>5=Will definitely pursue | Sum of Bivariate Career Interests (0-20)<br><br>Each Career Path:<br>a) If equal or below 3 = less interested = 0<br>b) If equal or above 4 = highly interested = 1 |
| The Familiarity with 20 Career Paths | NA | Which statement best describes your familiarity with the 20 career paths from the my Individual Development Plan (myIDP) shown in the Career Path Table | 1=I am not familiar with any of these career paths<br>2=I am familiar with a few of these career paths (between 1 and 6)<br>3=I am familiar with some of these career paths (between 7 and 12) | |

| | | | | |
|---|---|---|---|---|
| | | | 4=I am familiar with most of these career paths (between 13 and 19)<br>5= I am familiar with all of these career paths | |
| Career Training Attained | N/A | I am getting the training I need for my desired career path(s) | 0=Not applicable<br>1=Strongly disagree<br>2=Disagree<br>3=Neutral<br>4=Agree<br>5=Strongly agree | |
| Departmental Career Goal Support | Numerical | I am encouraged by my graduate program/department to pursue my career goals | 0=Not applicable<br>1=Strongly disagree<br>2=Disagree<br>3=Neutral<br>4=Agree<br>5=Strongly agree | |
| PI Encouragement | Yes | I am encouraged by my PI/thesis advisor to pursue career development activities toward my career goals<br>I am encouraged by my PI/thesis advisor to pursue my career goals | 0=I do not know<br>1=Strongly disagree<br>2=Disagree<br>3=Neutral<br>4=Agree<br>5=Strongly agree | |
| Citizenship | Yes | What is your citizenship status? (optional) | 1=US citizen since birth<br>2=Naturalized US citizen<br>3=Non-US citizen with permanent resident visa (green card)<br>4=Non-US citizen with temporary US visa | US = United States of America<br><br>Citizen = 1 or 2<br>Non-Citizen = 3 or 4 |
| Statistical Terms | N/A | | | ANOVA= analysis of variance<br>Tukey Corrections = for multiple comparisons |
| Other Terms | N/A | | | IRB=institutional review board<br>NIH BEST=National Institutes of Health Broadening Experiences in Scientific Training award<br>STEM=Science Technology Engineering and Math |

**S2 Table**

**Supplemental Table 2.** Inter-reliability of Items (α = 0.86)

| Deleted Variable | Item Total Correlation | Alpha (Item Deleted) |
|---|---|---|
| Item 1. Self-Assess abilities to pursue desired… | 0.71 | 0.83 |
| Item 2. Determine the steps to pursue desired… | 0.74 | 0.82 |
| Item 3. Seek advice from professionals in desired… | 0.65 | 0.84 |
| Item 4. Identify potential employers/institution… | 0.66 | 0.84 |
| Item 5. Achieve career goals | 0.67 | 0.84 |

**S3 Table 3. Full model ANOVA**

**Supplemental Table 3. 4-WAY ANOVA (Self-Efficacy with Gender x Citizenship x PI Career x Seniority)**

| **ANALYSIS OF VARIANCE** | | |
|---|---|---|
| Variable(s) | F-test | p-value |
| **Citizenship** | **50.784** | **.000** |
| **Gender** | **42.873** | **.000** |
| PI Career | **302.833** | **.000** |
| Seniority | 2.020 | .155 |
| **Citizenship x Gender** | **7.442** | **.006** |
| Citizenship x PI Career | .002 | .968 |
| Citizenship x Seniority | .858 | .354 |
| Gender x PI Career | .388 | .533 |
| Gender x Seniority | 2.587 | .108 |
| PI Career x Seniority | .637 | .425 |
| Citizenship x Gender x PI Career | .083 | .773 |
| Citizenship x Gender x Seniority | 1.639 | .201 |
| Citizenship x PI Career x Seniority | .114 | .736 |
| Gender x PI Career x Seniority | .255 | .614 |
| Citizenship x Gender x PI Career x Seniority | .085 | .771 |

**S4 Table 4. Contrasts (with Tukey corrections)**

| Tukey's multiple comparisons test | Significance | p-value |
|---|---|---|
| Male vs. Female | **** | <0.0001 |
| Male vs. Citizen Male Non-PI | **** | <0.0001 |
| Male vs. Citizen Male PI | **** | <0.0001 |
| Male vs. Citizen Female Non-PI | **** | <0.0001 |
| Male vs. Citizen Female PI | * | 0.011 |
| Male vs. Non-Citizen Male Non-PI | **** | <0.0001 |
| Male vs. Non-Citizen Male PI | * | 0.014 |
| Male vs. Non-Citizen Female Non-PI | **** | <0.0001 |
| Female vs. Citizen Male PI | **** | <0.0001 |
| Female vs. Citizen Female Non-PI | * | 0.0299 |
| Female vs. Citizen Female PI | **** | <0.0001 |
| Female vs. Non-Citizen Male PI | **** | <0.0001 |
| Female vs. Non-Citizen Female Non-PI | **** | <0.0001 |
| Citizen Male Non-PI vs. Citizen Male PI | **** | <0.0001 |
| Citizen Male Non-PI vs. Citizen Female PI | **** | <0.0001 |
| Citizen Male Non-PI vs. Non-Citizen Male PI | **** | <0.0001 |
| Citizen Male Non-PI vs. Non-Citizen Female Non-PI | **** | <0.0001 |
| Citizen Male PI vs. Citizen Female Non-PI | **** | <0.0001 |
| Citizen Male PI vs. Non-Citizen Male Non-PI | **** | <0.0001 |
| Citizen Male PI vs. Non-Citizen Female Non-PI | **** | <0.0001 |
| Citizen Male PI vs. Non-Citizen Female PI | **** | <0.0001 |
| Citizen Female Non-PI vs. Citizen Female PI | **** | <0.0001 |
| Citizen Female Non-PI vs. Non-Citizen Male PI | **** | <0.0001 |

| | | |
|---|---|---|
| Citizen Female Non-PI vs. Non-Citizen Female Non-PI | **** | <0.0001 |
| Citizen Female Non-PI vs. Non-Citizen Female PI | ** | 0.005 |
| Citizen Female PI vs. Non-Citizen Male Non-PI | **** | <0.0001 |
| Citizen Female PI vs. Non-Citizen Female Non-PI | **** | <0.0001 |
| Citizen Female PI vs. Non-Citizen Female PI | **** | <0.0001 |
| Non-Citizen Male Non-PI vs. Non-Citizen Male PI | **** | <0.0001 |
| Non-Citizen Male Non-PI vs. Non-Citizen Female Non-PI | * | 0.033 |
| Non-Citizen Male Non-PI vs. Non-Citizen Female PI | * | 0.031 |
| Non-Citizen Male PI vs. Non-Citizen Female Non-PI | **** | <0.0001 |
| Non-Citizen Male PI vs. Non-Citizen Female PI | **** | <0.0001 |
| Non-Citizen Female Non-PI vs. Non-Citizen Female PI | **** | <0.0001 |

**Supplemental Table 4 Legend.** Tukey-corrected P-values for each significant contrast are listed exactly, accompanied by asterisks such that *$p<.05$, **$p<.01$, ***$p<.001$, ****$p<.0001$.

### S5 Supplemental Figures

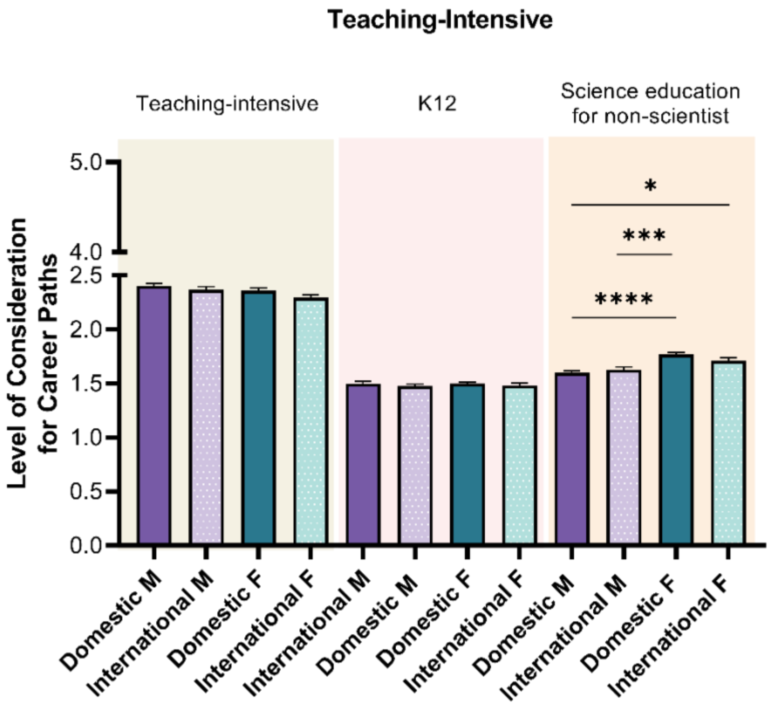


**S5 Figure 1: Level of consideration for Teaching-Intensive career paths.** Post-hoc t-tests were conducted between all possible pairings within each career path, as illustrated by each end of the respective bracket. P-values indicate significance of Tukey's multiple comparison tests, ****p<0.0001, ***p<0.001, and *p<0.05. *Note:* M=Male, F=Female. Color differences (purple and green) indicate the main variable gender, where purple = male, green = female; pattern indicates citizenship (no pattern = citizen; pattern = non-citizen).

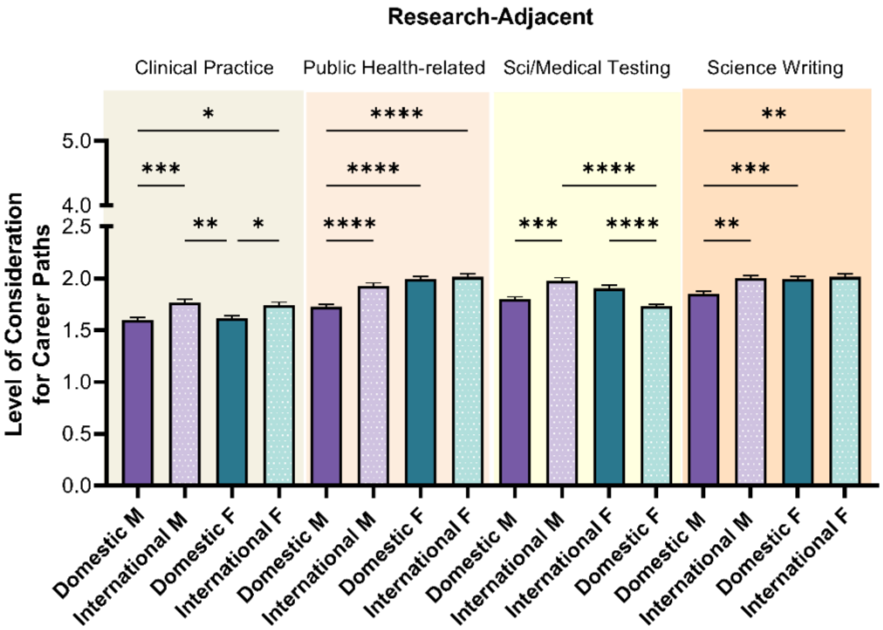


**S5 Figure 2: Level of consideration for Research-Adjacent career paths.** Post-hoc t-tests were conducted between all possible pairings within each career path, as illustrated by each end of the respective bracket. P-values indicate significance of Tukey's multiple comparison tests, ****p<0.0001, ***p<0.001, **p<0.01, and *p<0.05. *Note:* M=Male, F=Female. Color differences (purple and green) indicate the main variable gender, where purple = male, green = female; pattern indicates citizenship (no pattern = citizen; pattern = non-citizen).

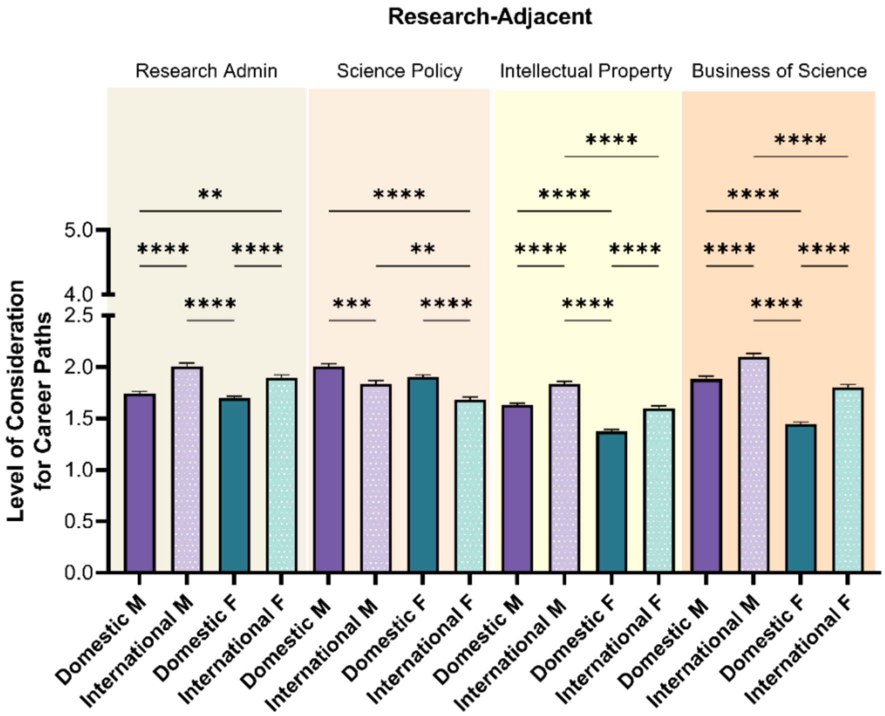


**S5 Figure 3: Level of consideration for other Research-Adjacent career paths.** Post-hoc t-tests were conducted between all possible pairings within each career path, as illustrated by each end of the respective bracket. P-values indicate significance of Tukey's multiple comparison tests, ****p<0.0001, ***p<0.001, and **p<0.01. *Note:* M=Male, F=Female. Color differences (purple and green) indicate the main variable gender, where purple = male, green = female; pattern indicates citizenship (no pattern = citizen; pattern = non-citizen).

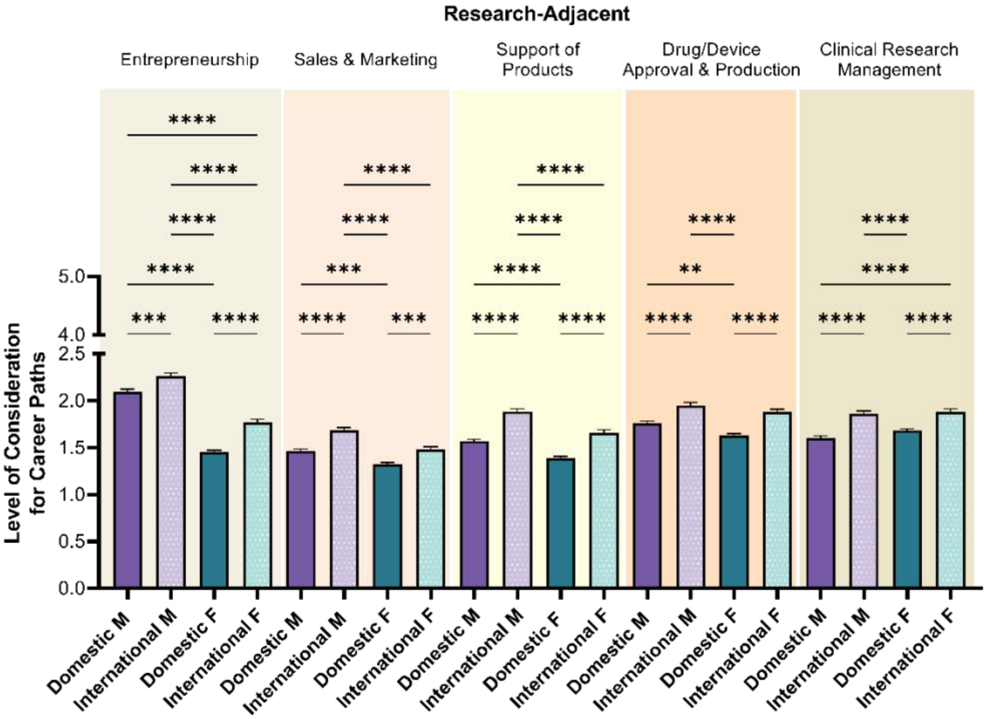


**S5 Figure 4: Level of consideration for Research-Adjacent career paths.** Post-hoc t-tests were conducted between all possible pairings within each career path, as illustrated by each end of the respective bracket. P-values indicate significance of Tukey's multiple comparison tests, ****p<0.0001, ***p<0.001, **p<0.01, and *p<0.05. *Note:* M=Male, F=Female. Color differences (purple and green) indicate the main variable gender, where purple = male, green = female; pattern indicates citizenship (no pattern = citizen; pattern = non-citizen).
